# Cold-induced suppression of myogenesis in skeletal muscle stem cells contributes to delayed muscle regeneration during hibernation

**DOI:** 10.1101/2025.05.30.654444

**Authors:** Tatsuya Miyaji, Ryuichi Kasuya, Mayuko Monden, Yutaka Tamura, Daisuke Tsukuamoto, Guangyuan Li, Shota Kawano, Yuri Watanabe, Yoshifumi Yamaguchi, Masatomo Watanabe, Mitsunori Miyazaki

## Abstract

Mammalian hibernators experience profound cold stress and prolonged physical inactivity during torpor periods; however, it is unclear how skeletal muscle stem cells (satellite cells; SCs) respond to these challenges. In this study, we demonstrate that SCs from a mammalian hibernator, Syrian hamster exhibit remarkable resistance to cold-induced cell death, which is associated with intrinsically higher expression of the antioxidant enzyme GPX4, which likely contributes to the suppression of ferroptosis. RNA-seq analysis revealed widespread downregulation of myogenesis-related genes following cold exposure, which suggests suppression of the myogenic program. Consistently, SCs exposed to cold stress exhibited reduced activation and differentiation capacities upon subsequent rewarming, with an increased number of quiescent Pax7-positive/MyoD-negative cells. Muscle regeneration was markedly delayed during hibernation, accompanied by decreased SC activation and macrophage infiltration, suggesting that cold-induced suppression of SC function underlies limited regenerative capacity in hibernating hamsters. Our results provide insight into the unique physiology of mammalian hibernators: SC viability is preserved, whereas regenerative activity is selectively suppressed during hibernation.

## Introduction

Hibernation is a survival strategy that organisms have developed throughout evolution to survive the harsh winter environment of cold ambient temperature (T_a_) and food deprivation. “Torpor” refers to the active reduction of energy metabolism and body temperature (T_b_) under conditions of cold and energy depletion in homeotherms that essentially maintain a constant T_b_^1–3^. Prolonged periods of torpor lasting for multiple days, occurring in a seasonal manner, are generally defined as hibernation^1,2,4,5^. There are more than 180 hibernating mammal species belonging to seven orders (*Monotremata*, *Marsupialia*, *Insectivora*, *Chiroptera*, *Primates*, *Rodentia*, and *Carnivora*), which indicates that approximately 5%–6% of all mammals use this survival strategy^1,2,6^. In small hibernators, such as hamsters and chipmunks, T_b_ decreases to near T_a_, often as low as 2°C–5°C during torpor^7,8^. The heart rate, respiratory rate, and whole-body energy metabolism are markedly reduced. These animals also experience rewarming of T_b_ during periodic arousal, thus repeating the torpor-arousal cycle every few days during hibernation^9–11^. Despite experiencing dramatic oscillations in T_b_, hibernating animals maintain physiological homeostasis without developing organ dysfunction^12^.

Living organisms are exposed to a variety of environmental stresses, including temperature changes. Excessive cellular stress can trigger cell death, leading to tissue damage and organ dysfunction. Cold-induced cell death (CICD), which is induced by prolonged exposure to hypothermic conditions, is mediated by ferroptosis^13^. Ferroptosis is a regulated form of cell death characterized by lipid peroxidation and iron dependency^14^. Syrian hamster-derived cultured cell lines exhibit resistance to CICD through the regulation of glutathione peroxidase 4 (GPX4), a glutathione-dependent antioxidant enzyme and a key regulator of ferroptosis^15,16^. In addition, primary hepatocytes from Syrian hamster are resistant to CICD^17^. Although such resistance may enable Syrian hamsters to survive extremely cold conditions during hibernation, it remains unclear whether other tissues or other hibernating species contain comparable mechanisms through ferroptosis suppression. Furthermore, the adaptive responses exhibited by surviving cells in hibernating animals following cold exposure remain poorly understood.

One remarkable feature of mammalian hibernators is their ability to preserve skeletal muscle mass despite experiencing prolonged physical inactivity^18^. Skeletal muscle is the largest organ in the body and plays an important role in force production and energy metabolism. Muscle loss due to aging or disuse has serious consequences for human health^19,20^. Satellite cells (SCs), the resident stem cells of skeletal muscle, are essential for muscle maintenance, repair, and regeneration^21^; however, it is unclear how SCs in mammalian hibernators adapt to cold exposure.

In this study, we isolated SCs from the skeletal muscles of mammalian hibernators and examined their response to cold exposure. Our findings were as follows: 1) SCs from the small hibernators, Syrian hamsters and chipmunks, exhibited resistance to CICD; 2) their proliferation was completely arrested under cold conditions, but resumed upon rewarming; 3) cold exposure downregulated the expression of genes associated with myogenic activation and differentiation; and 4) cold-induced suppression of myogenic processes was observed *in vivo* during hibernation following experimentally induced muscle injury.

## Results

### Isolation and characterization of satellite cells from hibernating mammals

To the best of our knowledge, SCs have not been previously isolated from mammalian hibernators, including the Syrian hamster. Using the method established by Yoshioka et al.^22^, we isolated SCs at high purity, as evidenced by the presence of Pax7- or MyoD-positive cells, which comprised over 95% of the primary cells isolated from hamster skeletal muscle (Supplemental Figure 1A–C). The SCs efficiently differentiated into myotubes when cultured in differentiation medium (DM), which confirmed their myogenic potential (Supplemental Figure 1D and E). Using the same method, we also successfully isolated SCs from chipmunks (Supplemental Figure 1F).

### Cold resistance and proliferative capacity of satellite cells from hibernating hamsters

We determined whether SCs from mammalian hibernators exhibited resistance to CICD. In SCs from mice, a nonhibernating species, cold treatment for 24 or 48 hours resulted in 61.7% and 85.4% propidium iodide (PI)-positive dead cells, respectively. In contrast, CICD was minimal (3.2% and 8.2%) in SCs from the Syrian hamster (Figure 1A–C). In an LDH cytotoxicity assay, mouse SCs exhibited over 80% cell death after 24 hours of cold exposure, whereas hamster SCs showed marked resistance (Figure 1D). CICD resistance was also observed in SCs isolated from chipmunk, which exhibited drastic T_b_ oscillations resulting from deep torpor and arousal during hibernation, like the hamster^23^. In contrast, marked CICD was observed in SCs from nonhibernating rats, as well as mice (Supplemental Figure 2A–C). These results suggest that CICD resistance may represent a unique cellular feature of mammalian hibernators that undergo profound reductions in T_b_ during torpor.

**Figure 1.**
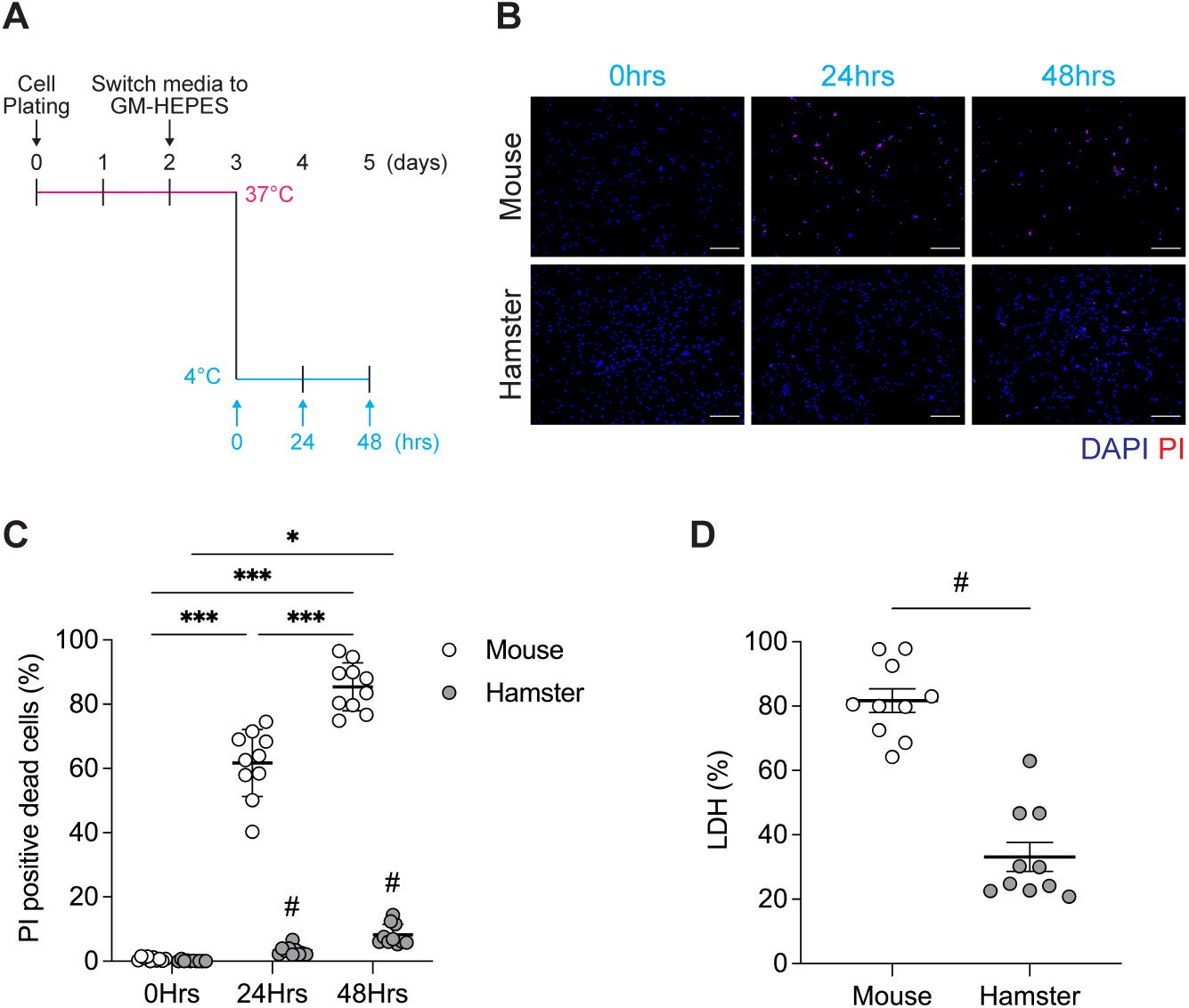
Satellite cells from hibernating hamsters exhibit resistance to cold-induced cell death. (A) Schematic overview of the experimental design to assess cold-induced cell death (CICD) in skeletal muscle satellite cells (SCs). SCs isolated from mice and Syrian hamsters were cultured at 37°C for 48 hours after plating, followed by a medium change to growth medium supplemented with HEPES (GM-HEPES) and an additional 24-hour incubation at 37°C. Cells were then subjected to cold exposure at 4°C for 24 or 48 hours without changing the medium. (B) Representative fluorescence images of SCs stained with propidium iodide (PI, red) and 4’, 6-diamidino-2-phenylindole (DAPI, blue) following cold exposure. PI-positive nuclei were frequently observed in mouse SCs, whereas hamster SCs largely retained viability. (C, D) Quantification of cell death by PI staining (C) and lactate dehydrogenase (LDH) release (D). In mouse SCs, both PI-positive cell counts and LDH release increased significantly over time. Hamster SCs exhibited minimal changes. Data are presented as mean ± SD (*n* = 10 per group). Statistical significance was assessed by two-way ANOVA followed by Tukey’s post hoc test. * p < 0.05, ** p < 0.01, *** p < 0.001 for time-dependent comparisons within species. # p < 0.05 for comparisons between mouse and hamster at the same time point. Scale bars: 200 μm (E–G).

To determine the role of ferroptosis in CICD, we assessed markers of cellular stress following cold exposure in mouse and hamster SCs. Hallmarks of ferroptosis, including increased reactive oxygen species (ROS) production (DCFH-DA staining) and the accumulation of intracellular free iron (FerroOrange), were observed in mouse SCs after 3–6 hours of cold exposure, whereas these responses were minimal in hamster SCs (Figure 2A, B). The level of GPX4 protein, an antioxidant enzyme that inhibits ferroptosis, was more abundant in hamster SCs compared with that in mouse SCs (Figure 2C, D). Neither the necroptosis inhibitor Necrostatin-1 nor the apoptosis inhibitor Z-VAD-FMK prevented CICD in mouse SCs, whereas the ferroptosis-specific inhibitor Ferrostatin-1 rescued cell viability (Figure 2E–G). These results indicate that CICD in nonhibernating mouse SCs is mediated by ferroptosis, whereas resistance in hamster SCs is attributed to the suppression of ferroptosis, which may be mediated by intrinsically higher GPX4 expression.

**Figure 2.**
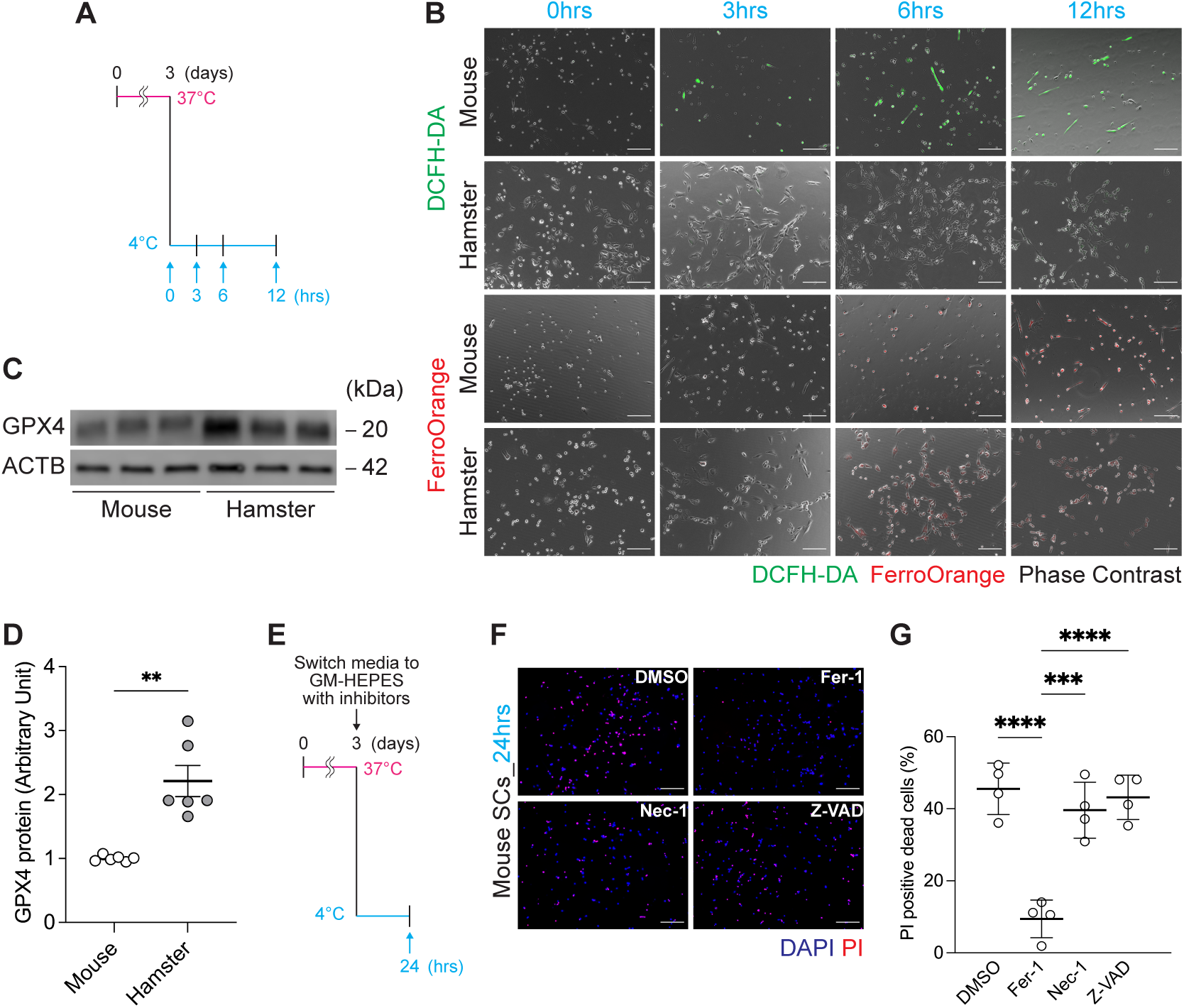
Cold-induced cell death in mouse satellite cells is mediated by ferroptosis and suppressed in hamster cells. (A) Schematic overview of the experimental design for evaluating oxidative stress and ferroptosis in satellite cells (SCs) exposed to cold. Mouse and hamster SCs were incubated at 4°C for 3, 6, or 12 hours, followed by staining with DCFH-DA and FerroOrange to detect reactive oxygen species (ROS) and intracellular free iron levels, respectively. (B) Representative fluorescence images of DCFH-DA and FerroOrange staining in mouse and hamster SCs after cold exposure. (C, D) Western blot analysis (C) and quantification (D) of glutathione peroxidase 4 (GPX4) and β-actin (ACTB) in mouse and hamster SCs under baseline conditions. *n* = 6 per group. Protein levels are expressed as arbitrary units. (E) Schematic overview of the experimental design for inhibitor assays. Mouse SCs were cultured at 37°C for 48 hours, followed by a 24-hour pre-incubation in GM-HEPES at 37°C. Cells were then subjected to 4°C cold exposure for 24 hours in the presence of vehicle (DMSO), ferroptosis inhibitor (Ferrostatin-1), necroptosis inhibitor (Necrostatin-1), or apoptosis inhibitor (Z-VAD-FMK). (F) Representative fluorescence images of PI and DAPI staining under each inhibitor condition. (G) Quantification of PI-positive cells in each group. Data are presented as mean ± SD (*n* = 4 per group). Statistical significance was assessed by one-way ANOVA followed by Tukey’s post hoc test. * p < 0.05, ** p < 0.01, *** p < 0.001, **** p < 0.0001 vs. DMSO-treated control. Scale bars: 200 μm (E–G).

To further examine the cellular response to cold stress, we pulsed the cells with the thymidine analog 5-ethynyl-2′ -deoxyuridine (EdU) to assess cell proliferation. After 24 hours of cold exposure, EdU-positive proliferating cells were absent in mouse and hamster SCs. Upon rewarming, mouse SCs exhibited minimal proliferation, whereas hamster SCs restored proliferation to levels comparable to that of preexposure conditions (Figure 3). These results indicate that hamster SCs resist CICD while retaining the capacity to proliferate following cold exposure and rewarming.

**Figure 3.**
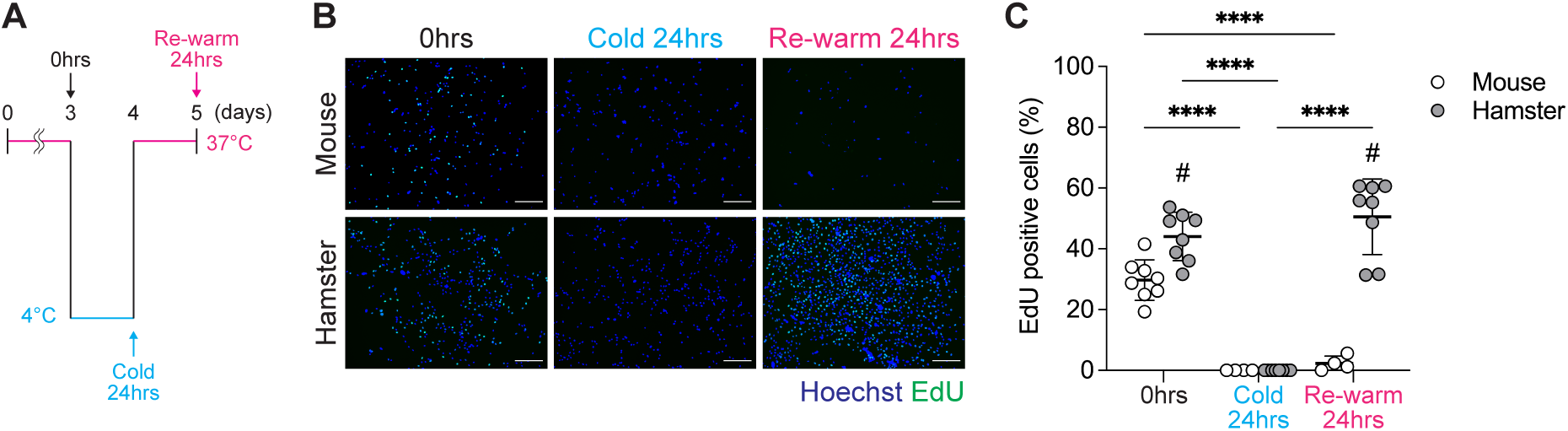
Satellite cells from hibernating hamsters resume proliferation after cold exposure and rewarming. (A) Schematic overview of the experimental design to assess satellite cell (SC) proliferative capacity after cold exposure and rewarming. SCs isolated from mice and Syrian hamsters were cultured at 37°C for 48 hours after plating, then incubated in GM-HEPES at 37°C for 24 hours, followed by cold exposure at 4°C for 24 hours. Cells were subsequently rewarmed and maintained at 37°C for another 24 hours. 5-ethynyl-2’-deoxyuridine (EdU) was added during the final 6 hours to label proliferating cells. (B) Representative fluorescence images of DAPI and EdU staining in mouse and hamster SCs under control, cold-exposed, and rewarming conditions. (C) Quantification of EdU-positive cells across conditions. Data are presented as mean ± SD (*n* = 4–8 per group). Statistical significance was assessed by two-way ANOVA followed by Tukey’s post hoc test. **** p < 0.0001 for comparisons between time points. # p < 0.05 for comparisons between mouse and hamster at the same time point. Scale bars: 200 μm (E–G).

### RNA-seq revealed cold-induced downregulation of myogenic genes in hamster SCs

RNA-seq analysis was performed to analyze the cellular responses to 24 hours of cold exposure in hamster SCs, and 64 differentially expressed genes (DEGs) were identified. DEGs were defined as those with a fold-change of |log2 FC| ≥ 1 (i.e., more >2-fold upregulation or less than half expression) and a Q-value <0.05. All 64 DEGs were downregulated in response to cold exposure (Figure 4A, B). Gene Ontology (GO) enrichment analysis (cellular component category) revealed significant enrichment for terms associated with striated muscle structures, including “Z-disc,” “striated muscle thin filament,” “myofibril,” “M band,” “myosin complex,” and “sarcolemma” (Figure 4C). The 64 DEGs included genes encoding structural components of muscle fibers (Myl4, Acta1, Myh3, Myl1, and Tnnt1), regulators of myogenic cell proliferation (Myom3^24^, Mstn^25^, Arx^26^, Egf^27^, St6gal2^28^, and Dtna^29^), and genes associated with myogenic differentiation (Aldh1a1^30^, Lmod3^31^, Hspb1^32,33^, Ripor2^34^, Casz1^35^, Kcnj2^36^, Abra^37^, Unc45b^38^, Mef2c^39^, Myoz2^40^, Smyd1^41^, Smpx^42^, and Mypn^43^) (Figure 4D–G). These results indicate that cold exposure induces broad transcriptional suppression of genes associated with muscle structure, SC activation, and myogenic differentiation in hamster SCs. This suggests a molecular mechanism that involves the downregulation of SC function under cold conditions.

**Figure 4.**
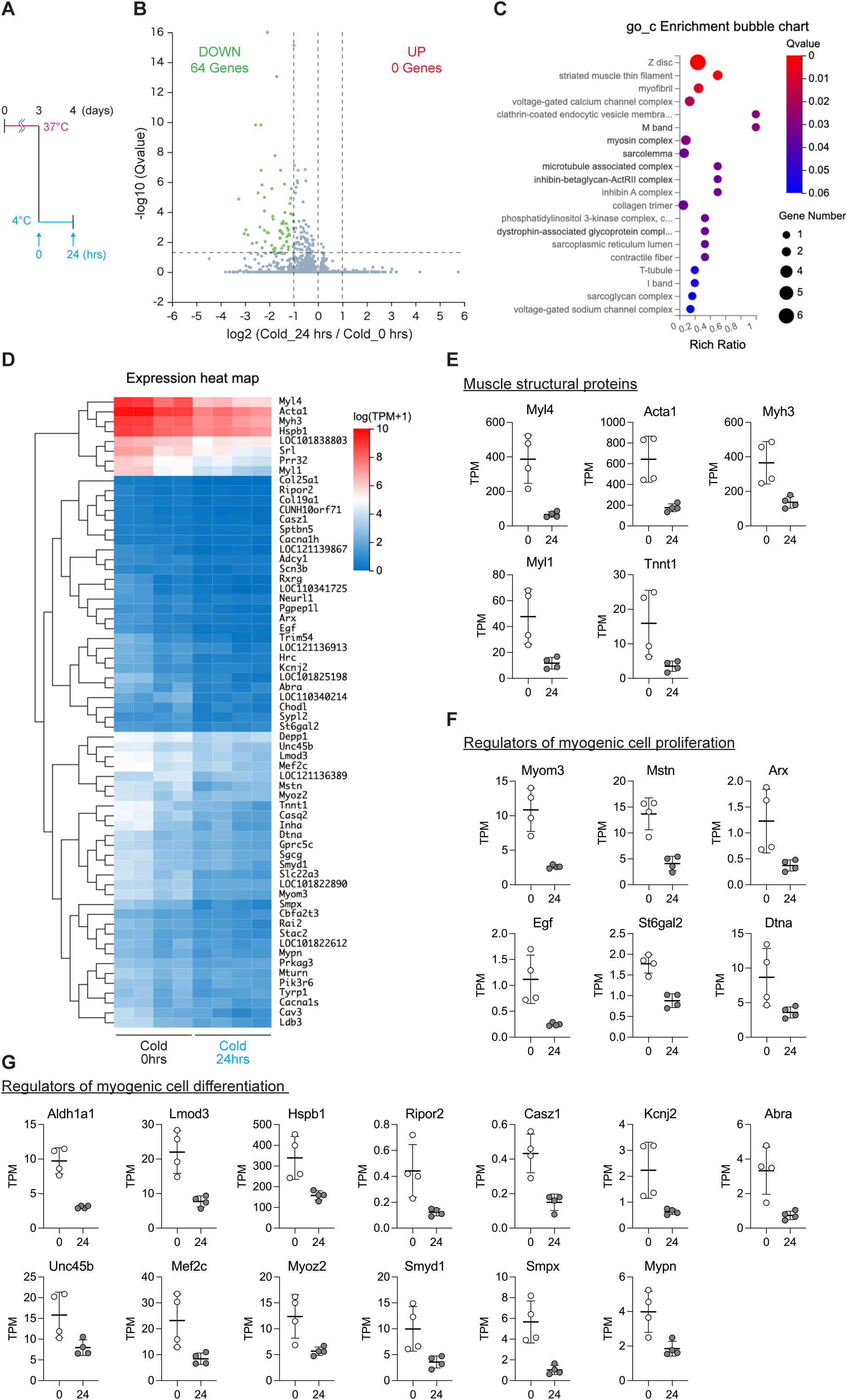
Cold exposure induces broad downregulation of myogenesis-related genes in hamster satellite cells. (A) Schematic overview of the experimental design for transcriptome analysis. Satellite cells (SCs) isolated from Syrian hamsters were cultured at 37°C for 48 hours after plating, followed by 24-hour incubation in GM-HEPES at 37°C, and then exposed to 4°C for 24 hours. Total RNA was extracted before and after cold exposure for RNA sequencing. (B) Volcano plot showing differentially expressed genes (DEGs) between control and cold-exposed SCs. A total of 64 genes were significantly downregulated after cold exposure (|log2 fold change| ≥ 1, Q < 0.05); no genes were upregulated. (C) Gene Ontology (GO) enrichment analysis (cellular component category) for the 64 downregulated DEGs. (D) Heatmap displaying expression levels of all 64 downregulated DEGs in control and cold-exposed SCs. (E–G) Selected subsets of the 64 downregulated DEGs, manually curated based on prior literature and known gene functions: (E) Genes encoding structural components of muscle fibers. (F) Genes involved in the regulation of satellite cell proliferation. (G) Genes associated with myogenic differentiation. RNA-seq was performed in biological triplicates (*n* = 4 per group). Differential expression was defined by |log2 fold change| ≥ 1 and Q < 0.05. For panels (E–G), gene expression levels are shown as transcript per million (TPM) values directly derived from RNA-seq output without further normalization.

### Cold exposure suppresses myogenic activation and differentiation in hamster SCs

To determine the effect of cold exposure on myogenic progression in CICD-resistant hamster SCs, we measured the expression of Pax7 and MyoD in cells that were rewarmed following 24 hours of cold treatment. Compared with the noncold-exposed controls, the proportion of MyoD-positive cells, a marker of SC activation, was significantly reduced in cold-exposed hamster SCs, whereas the proportion of Pax7-positive/MyoD-negative (Pax7⁺/MyoD⁻) quiescent satellite cells was concomitantly increased (Figure 5A–C). In addition, the percentage of myogenin-positive cells, a marker of myogenic differentiation, was significantly decreased following cold exposure in hamster SCs subjected to differentiation induction (Figure 5D–F). Furthermore, despite being subjected to the same duration of differentiation induction, both the protein expression of the myosin heavy chain, a hallmark of late-stage differentiation, and the fusion index were reduced, indicating the inhibition of the myogenic process in response to cold exposure (Figure 5G–L).

**Figure 5.**
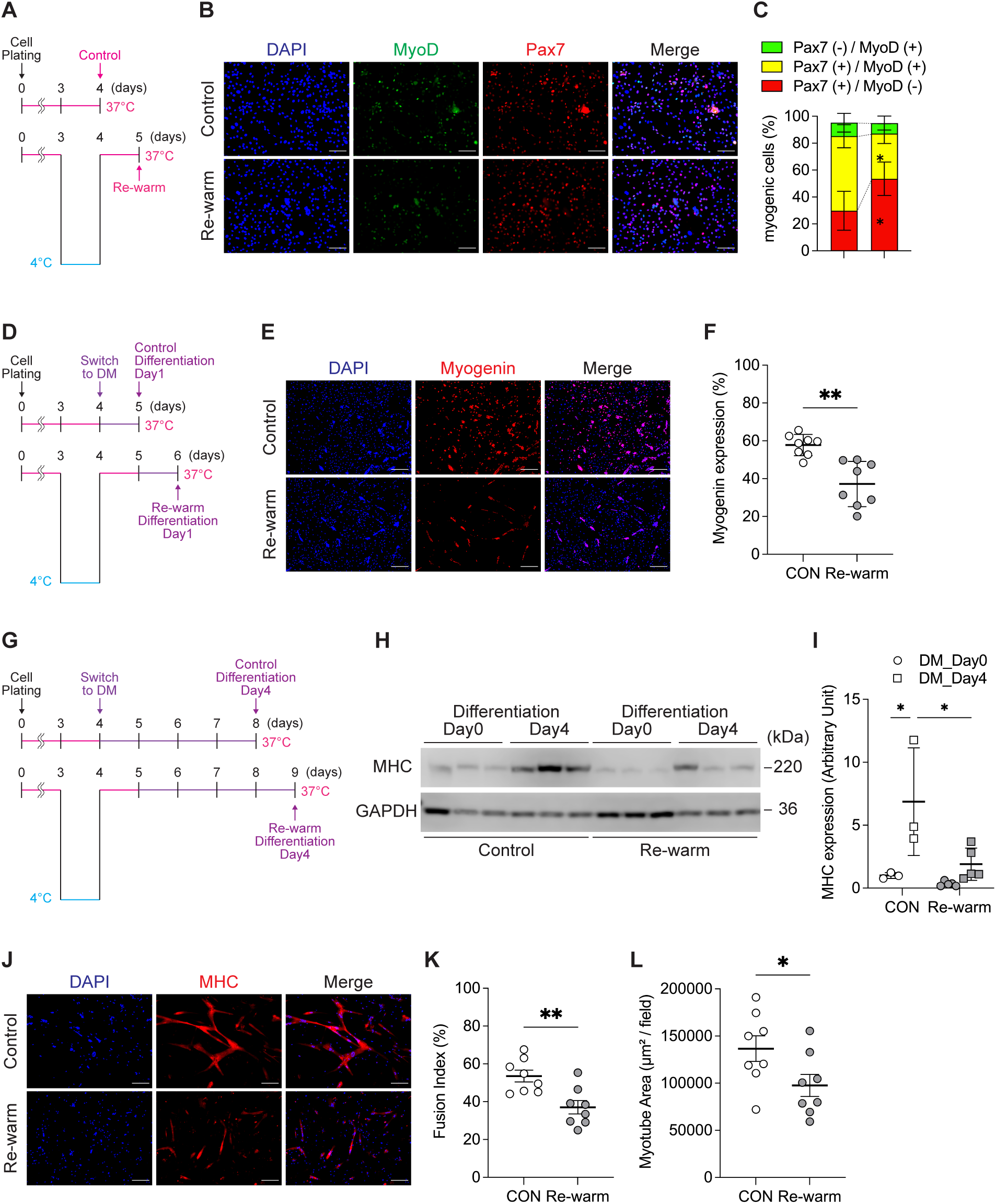
Cold exposure suppresses satellite cell activation and differentiation in hamster cells. (A) Schematic overview of the experimental design for cold exposure and rewarming to evaluate satellite cell activation. Hamster satellite cells (SCs) were cultured at 37°C for 48 hours after plating, then incubated in GM-HEPES at 37°C for 24 hours, followed by 24-hour exposure to 4°C. Cells were then rewarmed and maintained at 37°C for 24 hours before analysis. (B) Representative fluorescence images of Pax7 and MyoD immunostaining in control and rewarmed SCs. (C) Quantification of the relative proportions of three cell populations—Pax7⁺/MyoD⁻, Pax7⁺/MyoD⁺, and Pax7⁻/MyoD⁺—within the total cell population. (D) Schematic overview of the experimental design for cold exposure, rewarming, and subsequent induction of differentiation. (E) Representative images of myogenin immunostaining in control and rewarmed SCs following 24 hours in differentiation medium. (F) Quantification of myogenin-positive cells after 24 hours of differentiation. (G) Schematic overview of the experimental design for cold exposure, rewarming, and subsequent induction of differentiation. During the differentiation period, cells were maintained in DM for 4 days, with the medium replaced with fresh DM every other day. (H) Western blot analysis of myosin heavy chain (MHC) and glyceraldehyde-3-phosphate dehydrogenase (GAPDH) protein levels at differentiation day 0 and day 4. GAPDH was used as a loading control. (I) Quantification of MHC protein band intensity. (J) Representative images of MHC immunostaining used for fusion index calculation. (K) Quantification of fusion index (percentage of nuclei within MHC-positive myotubes). (L) Quantification of the total area of MHC-positive regions. Statistical analyses were performed as follows: (F), (K), and (L), unpaired two-tailed Student’s t-test; (I), two-way ANOVA followed by Tukey’s post hoc test. Sample sizes: (F), (K), and (L), *n* = 8 per group; (I), *n* = 3–5 per group. Data are presented as mean ± SD. * p < 0.05, ** p < 0.01, *** p < 0.001; significance levels correspond to comparisons indicated in each panel. # p < 0.05 for comparisons between control and rewarmed groups at the same time point. Scale bars: 200 μm (E–G).

### Hibernation delays skeletal muscle regeneration

Having established that cold exposure of primary cultured SCs from Syrian hamsters suppresses myogenic progression, we determined the effect of hibernation on skeletal muscle regeneration *in vivo*. Syrian hamsters were transferred from a standard environment (ambient temperature: 23°C ± 1°C, light 14 h, dark 10 h) to a winter-like condition (ambient temperature: 5°C, light 8 h, dark 16 h), where they entered hibernation over a period of 2 to 4 months. During hibernation, the animals repeatedly cycled between states of deep torpor, marked by a pronounced reduction in T_b_, and periodic arousal, during which T_b_ returned to normothermic levels (Figure 6A).

**Figure 6.**
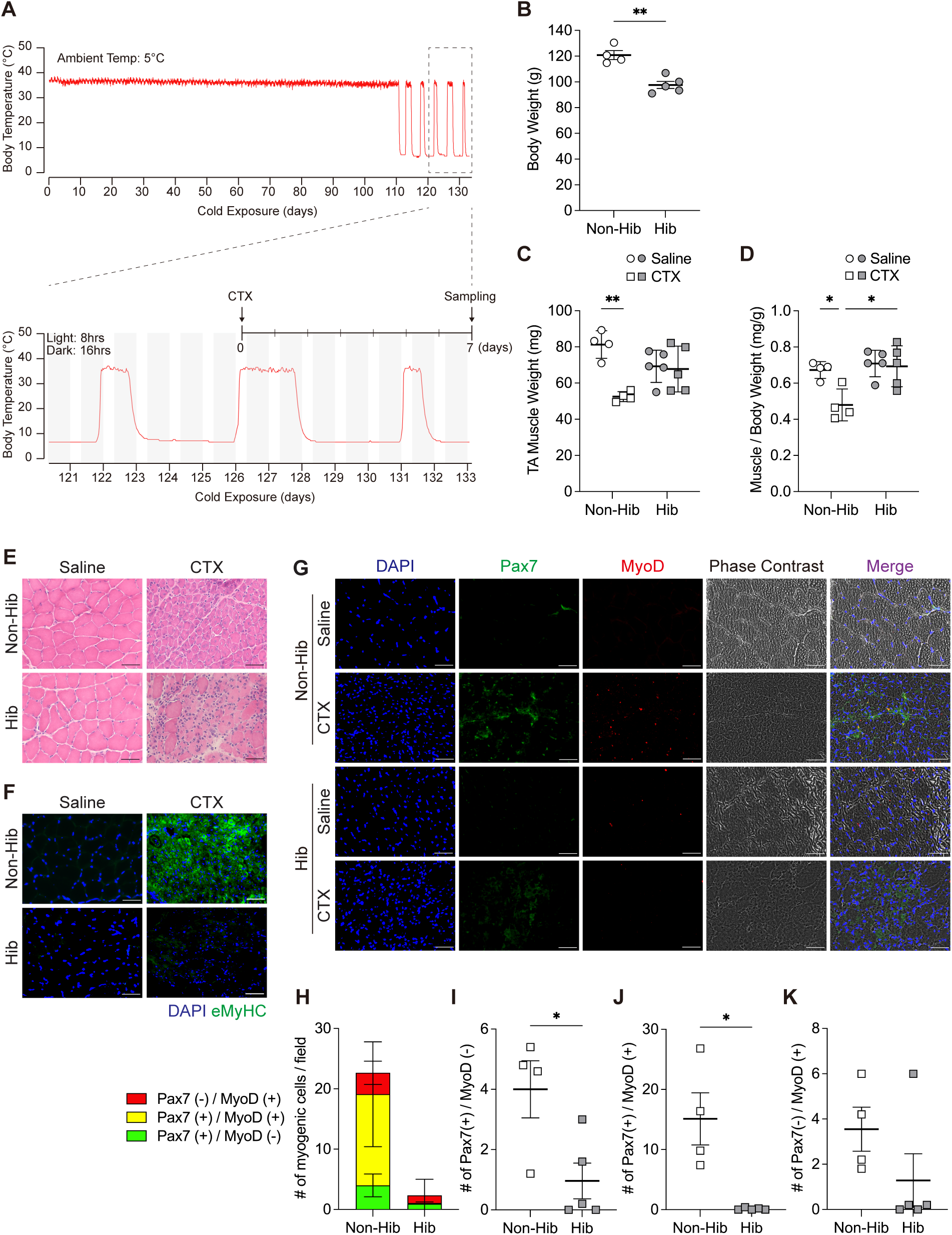
Skeletal muscle regeneration is suppressed during hibernation in Syrian hamsters. (A) Representative body temperature fluctuations recorded in a Syrian hamster over the course of hibernation. The period corresponding to hibernation is enlarged to indicate the timing of cardiotoxin (CTX) injection during an interbout arousal and tissue collection 7 days later during deep torpor. (B) Body weight in nonhibernating (Non-Hib) and hibernating (Hib) animals at the time of tissue collection. (C) Absolute weights of the tibialis anterior (TA) muscles, with saline injected into one leg and CTX into the contralateral side. (D) TA muscle weights normalized to body weight. (E) Representative hematoxylin and eosin staining of TA cross sections. (F) Representative immunofluorescence images of embryonic myosin heavy chain expression. (G) Representative images showing Pax7 (green) and MyoD (red) expression, along with DAPI staining and phase contrast. Merged images are also shown. (H) Total number of Pax7⁺ and/or MyoD⁺ cells per field of view. For each sample, at least five random fields were imaged, and the average number of positively stained cells per field was calculated and used as one biological replicate. (I–K) Separate quantification of Pax7⁺/MyoD⁻ (I), Pax7⁺/MyoD⁺ (J), and Pax7⁻/MyoD⁺ (K) cell populations in Non-Hib and Hib groups. Statistical analysis: (B, I–K), unpaired two-tailed Student’s t-test; (C, D), two-way ANOVA followed by Tukey’s post hoc test. Sample sizes: Non-Hib, *n* = 4; Hib, *n* = 5. Data are presented as mean ± SD. * p < 0.05, ** p < 0.01. Scale bars: 50 μm (E–G).

Compared with the nonhibernating controls housed in the same cold and short-day conditions, hibernating hamsters exhibited reduced body weight (Figure 6B). Following cardiotoxin (CTX) injection into the tibialis anterior (TA) muscle, nonhibernating controls showed a significant reduction in muscle weight compared with the contralateral saline-injected side, whereas muscle weight was maintained in the hamsters during hibernation (Figure 6C, D). Histological analysis at day 7 postinjury revealed numerous regenerating myofibers with small diameters and central nuclei in the nonhibernating group, whereas the hibernating group exhibited limited fiber regeneration, with persistent cellular presence in the interstitial space, which suggested delayed or arrested regenerative progression (Figure 6E). The expression of embryonic myosin heavy chain (eMyHC), a marker of regenerating fibers, was observed in the nonhibernating group, but absent in hamsters during hibernation (Figure 6F). Furthermore, the proportion of Pax7- or MyoD-positive SCs was significantly increased during regeneration in nonhibernating controls; however, this enhancement was markedly suppressed during hibernation (Figure 6G–K). These *in vivo* findings demonstrate that hibernation, characterized by cycles of deep torpor and arousal, suppresses the regenerative activation of SCs and delays muscle regeneration following injury.

### Attenuated inflammatory response during muscle regeneration in hibernating animals

Regulation of the tissue microenvironment, particularly immune cell infiltration, has an important role in muscle regeneration^44^. In nonhibernating controls, the number of CD68-positive macrophages was significantly increased in the damaged muscle, whereas macrophage infiltration was markedly reduced in the animals during hibernation. Both proinflammatory M1-like macrophages (CD68-positive/iNOS-positive) and anti-inflammatory M2-like macrophages (CD68-positive/CD206-positive) were sparsely detected in the injured muscles of the hibernating group (Figure 7). The results suggest that hibernation suppresses both SC activation and myogenic regeneration, while simultaneously attenuating the inflammatory response, which is essential for effective muscle repair.

**Figure 7.**
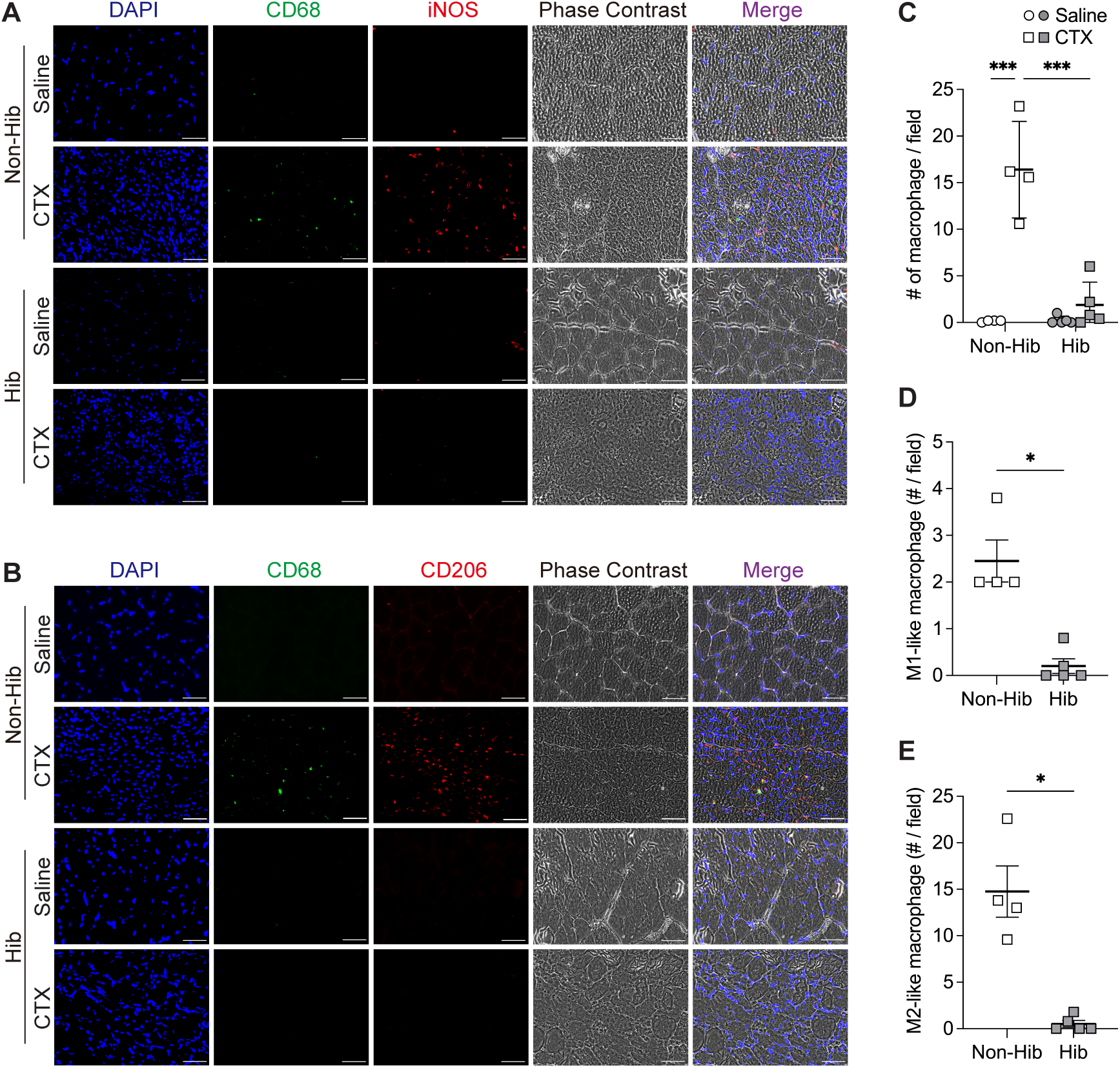
Macrophage infiltration is suppressed during hibernation in regenerating muscle. (A, B) Representative immunofluorescence images stained for CD68 (pan-macrophage marker), iNOS (M1 marker; panel A), or CD206 (M2 marker; panel B), along with DAPI and phase contrast. (C) Quantification of total macrophages defined as CD68⁺/DAPI⁺ cells in both saline- and CTX-injected muscles. (D) Quantification of M1-like macrophages defined as CD68⁺/iNOS⁺/DAPI⁺ cells. (E) Quantification of M2-like macrophages defined as CD68⁺/CD206⁺/DAPI⁺ cells. For quantification, at least five randomly selected fields were imaged per sample. The number of marker-positive cells per field was averaged and used as one biological replicate. Statistical analysis: (C), two-way ANOVA followed by Tukey’s post hoc test; (D, E), unpaired two-tailed Student’s t-test, performed only on CTX-injected samples due to undetectable marker expression in saline-injected muscles. Sample sizes: Non-Hib, *n* = 4; Hib, *n* = 5. Data are presented as mean ± SD. * p < 0.05, *** p < 0.001. Scale bars: 50 μm (A, B).

## Discussion

In the present study, we demonstrated that skeletal muscle SCs from hibernating animals exhibit marked resistance to CICD by suppressing ferroptosis, an iron-dependent form of regulated cell death. This intrinsic cellular process likely represents a mechanism that enables hibernating animals to survive periods of low T_b_ associated with hibernation under cold temperatures. In contrast, cold exposure significantly suppressed SC activation and differentiation *in vitro* and delayed muscle regeneration *in vivo*. Moreover, macrophage infiltration into the damaged muscle tissue was markedly reduced during hibernation. These results suggest that during hibernation, a state characterized by a profound reduction in T_b_ and multiple acute cellular responses, including myogenesis and inflammation, are concurrently downregulated.

Previous studies showed that culturing muscle SCs or myoblasts at subphysiological temperatures (e.g., 30°C–37°C) delays proliferation and differentiation, potentially leading to the formation of smaller myofibers^45–47^. Although our findings are generally consistent with temperature-dependent suppression of myogenesis, the culture conditions used in the present study (exposure to 4°C) represent a far more extreme cold challenge. Under these conditions, SC proliferation was completely arrested, yet the proliferative capacity was restored upon rewarming. This reversible suppression highlights a unique aspect of SC behavior in hibernating animals, in which cold exposure imposes a temporary block on proliferation without inducing irreversible cell cycle arrest. Interestingly, this occurred despite the marked suppression of canonical myogenic regulators, such as MyoD. This suggests that SC proliferation can occur independently of MyoD expression, which is consistent with previous reports indicating that MyoD-negative SCs are capable of division under certain conditions, such as mechanical overload, in which sustained HeyL expression suppresses MyoD while promoting SC expansion^48^.

Mechanistically, resistance to CICD in SCs from hibernators may be attributed, in part, to the intrinsically higher expression of GPX4, a key regulator of ferroptosis. GPX4 is an antioxidant enzyme that prevents lipid peroxidation. Its elevated expression in SCs from hibernating animals likely contributes to the suppression of ferroptotic cell death during cold exposure. This mechanism may represent an important aspect of cellular resilience that allows SCs to remain viable under extreme cold stress^15,16^.

We also observed a concomitant cold-induced increase in the proportion of Pax7⁺/MyoD⁻ SCs, suggesting the preservation of the quiescent SC pool. Maintaining quiescence is important for long-term muscle homeostasis and regenerative capacity^49^. Therefore, the suppression of SC activation during cold exposure may represent a protective strategy that safeguards the stem cell reserve during prolonged environmental stress, thus enabling future regeneration once favorable conditions return.

Our *in vivo* experiments demonstrated that muscle regeneration was delayed during hibernation, which is consistent with previous studies in 13-lined ground squirrels^50^. In this model, delayed muscle regeneration during hibernation was associated with reduced expression of transforming growth factor beta 1, a key profibrotic factor in skeletal muscle, along with minimal fibrosis. This suggests that a slower regenerative response may contribute to the preservation of muscle integrity, while avoiding pathological remodeling. Our results extend these observations by demonstrating that the suppression of muscle regeneration in hibernating animals is accompanied by reduced macrophage infiltration following injury. Macrophages play an important role in promoting muscle regeneration through metabolic and signaling interactions with SCs^51^. Thus, the attenuation of regenerative and inflammatory responses during hibernation likely reflects physiological adaptation to reduce the metabolic expenditure associated with tissue remodeling. The physiological significance of these adaptations likely resides in the minimization of energy expenditure during periods when energy is limited. During hibernation, the suppression of metabolically demanding processes may represent a physiological adaptation that conserves energy and supports the maintenance of tissue integrity during cold and nutrient-poor conditions.

Although our RNA-seq analysis revealed the downregulation of myogenesis-related genes following cold exposure, the specific molecular pathways mediating these transcriptional changes remain to be identified. Future studies examining the epigenetic mechanisms, chromatin remodeling, and transcriptional regulation underlying cold-induced gene suppression will be important to determine how cold exposure alters the myogenic program in SCs and may ultimately provide insight into the adaptive regulation of SC function in hibernating animals. Elucidating these mechanisms may also inform future studies for tissue preservation or regenerative interventions in settings, such as therapeutic hypothermia and spaceflight-induced muscle atrophy.

## Materials and Methods

All reagents, antibodies, and commercially available materials used in this study are listed in Supplemental Table 1.

### Animals

All animal procedures were approved by the Animal Ethics and Research Committees of Hiroshima University (No. A22-7-6 and No. A24-38-2), Fukuyama University (No. 2022-A-12 and 2025-A-5), and Kitasato University (No. SA2402), and complied with the institutional guidelines. The hamsters (Slc:Syrian), mice (C57BL/6JmsSlc), and rats (Slc:SD) used for SC isolation were sexually mature (8–16 weeks old) males. They were purchased from Japan SLC (Shizuoka, Japan) and maintained in a temperature- and humidity-controlled animal facility (23°C ± 1°C, 50%–60% humidity) at Hiroshima University under a 12 h light/12 h dark cycle. They were provided autoclaved water and a certified rodent diet *ad libitum*. Skeletal muscle samples were harvested under anesthesia followed by euthanasia (mice: cervical dislocation under 2.0% isoflurane; rats and hamsters: intraperitoneal overdose of sodium pentobarbital at 200 mg/kg).

For the hibernation experiments, Syrian hamsters were bred at Fukuyama University using breeding pairs (Slc:Syrian) obtained from Japan SLC (Shizuoka, Japan). All of the animals were male and housed under standard conditions (23°C ± 1°C, 14 h light/10 h dark cycle) until 8 weeks of age, at which time they were intraperitoneally implanted with a temperature logger. Following a 1-week recovery period, they were transferred to a controlled environment (5°C, 8 h light/16 h dark) designed to simulate a winter-like environment to induce hibernation. Animals were individually housed with *ad libitum* access to diets (Labo MR Standard, Nihon Nosan, Japan). The duration required for entry into hibernation varied among individuals.

The chipmunks (*Tamias sibiricus*) used for skeletal muscle collection were approximately 6 months old males, purchased from Pure Animal (Tokyo, Japan). They were individually housed at 23℃ under a 12 h light/12 h dark cycle in the animal facility at Kitasato University, with free access to a standard rodent diet and water. Skeletal muscle samples were collected from non-hibernating individuals that were euthanized under deep isoflurane anesthesia.

### Satellite cell isolation and culture

SCs were isolated according to the method of Yoshioka et al.^22^ Briefly, hindlimb skeletal muscles were dissected, minced with sterile surgical scissors, and digested in 0.2% type II collagenase solution at 37°C for 60 min. The tissue was gently homogenized using a 20-G syringe needle and filtered through a 40 µm nylon mesh strainer. The filtrate was centrifuged at 500 × g for 5 min and the pellet was resuspended in GM consisting of DMEM supplemented with 30% fetal bovine serum, 1% chick embryo extract, 10 ng/mL basic fibroblast growth factor, and 1× penicillin-streptomycin solution. The SCs were purified using a classical preplating method, followed by sequential replating to remove fibroblasts and enrich SCs. Purified SCs were plated onto Matrigel-coated culture dishes and maintained at 37°C in GM in a standard 5% CO_2_ incubator. The medium was changed every other day. To induce differentiation, GM was replaced with DM consisting of DMEM supplemented with 2% horse serum and 1× penicillin-streptomycin. Differentiation was induced after 3 days of culture in GM, and the cells were maintained in DM for up to 4 days depending on the experiment.

### Cold exposure and rewarming

To induce cold stress, SCs were precultured at 37°C for 48 h, followed by 24 h in GM supplemented with 100 mM HEPES (pH 7.4) (GM-HEPES) at 37°C. HEPES was added to stabilize the pH during cold exposure and prevent temperature-dependent pH fluctuations. Sealed plastic containers containing a small amount of water were preequilibrated in a conventional refrigerator maintained at 4°C and monitored with a thermometer. The SC culture dishes were placed into these containers and stored at 4°C for 24–48 h without medium change. For rewarming, the SCs were returned to 37°C for 24 h prior to analyses.

### Cell death and cytotoxicity assays

SCs were incubated with PI (5 µg/mL in PBS) for 5 min at room temperature under dark conditions, fixed with 4% paraformaldehyde (PFA) in PBS for 10 min, and counterstained with 4’,6-diamidino-2-phenylindole (DAPI). Fluorescence images were obtained using a BZ-X800 microscope system. LDH activity in the culture medium was measured using an LDH Cytotoxicity Assay Kit based on the manufacturer’s instructions with minor modifications. A blank control consisting of medium without cells was used to determine the background absorbance. The absorbance at 490 nm was measured using a Multiskan GO microplate reader.

### Reactive oxygen species and iron imaging

Cold-induced mitochondrial oxidative stress was assessed using a ROS Assay Kit (Highly Sensitive DCFH-DA). FerroOrange was used to detect intracellular ferrous iron (Fe²⁺) as a ferroptosis marker. Cells were stained with DCFH-DA or 1 µM FerroOrange based on the manufacturer’s instructions and imaged using a BZ-X800 fluorescence microscope system.

### Inhibitor assays

To analyze cell death pathways during cold exposure, the following inhibitors were added: 1 µM Ferrostatin-1 (ferroptosis), 20 µM Necrostatin-1 (necroptosis), and 20 µM Z-VAD-FMK (apoptosis). DMSO was used as a vehicle control. Inhibitor-containing GM-HEPES was used throughout the 24-h cold treatment.

### Western blot analysis

The cell samples were lysed in a 1× concentration of sample buffer and collected using a cell scraper. The lysates were triturated through a 29G syringe needle and heated at 95°C for 5 min. Equal volumes of lysates were separated using a precast polyacrylamide gel system and transferred to PVDF membranes. The membranes were blocked and then incubated with appropriate dilutions of primary and secondary antibodies. ACTB and GAPDH detection served as a loading control to confirm equivalent loading. Detection was done using chemiluminescence reagents, and images were acquired using a C-DiGit Blot Scanner with Image Studio Digits 5.2 software.

### EdU incorporation assay

Cell proliferation was measured using the Click-iT EdU Cell Proliferation Kit for Imaging and Alexa Fluor 488 dye, following the manufacturer’s protocol. EdU (10 µM) was added during the final 6 h of each experimental condition (control, cold exposure at 4°C, or rewarming at 37°C), while maintaining the respective environmental settings. The cells were fixed in 4% PFA for 15 min at room temperature and permeabilized with 0.5% Triton X-100 in PBS for 20 min. The Click-iT reaction cocktail was added and incubated in the dark at room temperature for 30 min. The cells were counterstained with Hoechst 33342 for 30 min in the dark and washed with PBS. Fluorescence images were acquired using a BZ-X800 microscope system, and the cells were counted using the Hybrid Cell Count module of the BZ-X Analyzer software, based on Hoechst and Alexa Fluor 488 signals in five randomly selected nonoverlapping fields.

### RNA sequencing and bioinformatics

Total RNA was extracted using the ISOGEN II and RNeasy Mini Kit. Total RNA quantitation and quality checks for purity and integrity were performed using a 5300 Fragment Analyzer System. RNA libraries and transcriptome sequencing were performed by BGI (Guangdong, China). Briefly, the libraries were prepared with the Optimal Dual-mode mRNA Library Prep Kit based on the manufacturer’s recommended protocol. Each library was then sequenced using the DNBSEQ-G400 platform with a paired-end sequence length of 150 bp (PE 150). DEGs were filtered based on a >2.0-fold increase or <1/2 decrease (|log2FC| ≥ 1) and a Q value <0.05. Bioinformatics analysis, including GO analysis and heat mapping, was performed using Dr. Tom software provided by BGI (Guangdong, China). The RNA-seq datasets generated during the current study were deposited in the Gene Expression Omnibus under accession code (GSE296910).

### Immunofluorescence staining of the cultured cells

Cells were fixed with 4% PFA, permeabilized with 0.3% Triton X-100, and blocked with 5% goat serum. Cell samples were then incubated overnight at 4°C with primary antibodies diluted at the appropriate concentrations. Secondary antibodies conjugated with Alexa Fluor 488 or 594 were used appropriately, considering the combination of host animal species and immunoglobulin subtypes of the primary antibodies. Cells were then treated with a DAPI solution for nuclear counterstaining. All images were obtained using a BZ-X800 microscope system (Keyence, Osaka, Japan).

### In vivo muscle regeneration

Regeneration of the skeletal muscle was induced by the intramuscular injection of CTX, which is a snake venom-derived toxin that causes acute myofiber damage. During the periodic arousal phase of hibernation, 100 µL of 10 µM CTX in PBS was injected into the TA muscle under isoflurane inhalation anesthesia. As a control, the contralateral TA muscle was injected with an equal volume of sterile saline. Skeletal muscle tissue was harvested 7 days postinjection during deep torpor following euthanasia and decapitation using an intraperitoneal administration of a medetomidine (0.75 mg/kg), midazolam (4 mg/kg), and butorphanol (5 mg/kg) mixture.

### Histology and immunostaining of muscle tissue

Skeletal muscle specimens were frozen in isopentane cooled with liquid nitrogen. They were sectioned into 10-µm-thick sections using a cryostat (Microm NX50H or Cryostat HM525NX, Thermo Fisher Scientific, Rockford, IL, USA), and stored at –80°C until further analysis. H&E staining involved fixing sections in 4% PFA and staining with Mayer’s hematoxylin solution for 5 min, followed by eosin staining for 5 min. For immunohistochemistry, the sections were fixed with 4% PFA, permeabilized with 0.1% Triton X-100, and blocked with 1% bovine serum albumin. The samples were incubated overnight at 4°C with primary antibodies diluted at the appropriate concentration. Secondary antibodies conjugated with fluorescence labeling were used appropriately, considering the combination of the host animal species and the immunoglobulin subtypes of the primary antibodies. The antibodies and combinations are listed in Supplementary Table 2. For the detection of M1-like and M2-like macrophages, a fluorescent dye-conjugated CD68 antibody and an antibody labeling kit for rabbit IgG (FlexAble 2.0 CoraLite Plus 555 Antibody Labeling Kit for Rabbit IgG) were used to accommodate the shared host species. The sections were mounted with DAPI Fluoromount-G for nuclear counterstaining. The images were obtained using a BZ-X800 microscope system (Keyence, Osaka, Japan). Five to six nonoverlapping areas were randomly selected and uniformly distributed across each muscle section for quantitative analysis.

### Quantification and statistical analysis

Quantitative analyses included cell viability, EdU incorporation, immunofluorescence signal intensity, muscle fiber cross-sectional area, and macrophage infiltration. At least five randomly selected, nonoverlapping fields per sample were analyzed using BZ-X Analyzer software with the Hybrid Cell Count module (Keyence). The average per sample was considered one biological replicate. Statistical tests were selected based on the experimental design and the number of groups. For comparisons between two groups, unpaired two-tailed Student’s t-tests were performed. For comparisons of more than two groups or involving two independent variables, one-way or two-way ANOVA followed by Tukey’s multiple comparison test, was used. All data are presented as the mean ± standard deviation. Significance thresholds were defined as p < 0.05, p < 0.01, p < 0.001, and p < 0.0001. Detailed statistical methods, n values, and group comparisons are described in the corresponding figure legends.

## Acknowlegements

We would like to thank Dr. Masamitsu Sone, Institute of Low Temperature Science, Hokkaido University, for valuable discussions and insightful comments on this study, and Dr. Yasuo Kitajima, Department of Immunology, Graduate School of Biomedical and Health Sciences, Hiroshima University, for technical guidance on the isolation of satellite cells from skeletal muscle tissue. The MYH1E antibody (MF 20) developed by Dr. Fischman, D.A. (Cornell University Medical College) was obtained from the Developmental Studies Hybridoma Bank, created by the NICHD of the NIH and maintained at The University of Iowa, Department of Biology, Iowa City, IA 52242.

## Author Contribution Statement

MM conceived and designed the project; TM, RK, MM, YT, DT, GL, SK, YW, and MW acquired, analyzed, and interpreted the data; TM, RK, YY, MW, and MM wrote, revised, and edited the paper. MM directed overall research project.

