## Supplemental_Figure_1 for "Cold-induced suppression of myogenesis in skeletal muscle stem cells contributes to delayed muscle regeneration during hibernation"

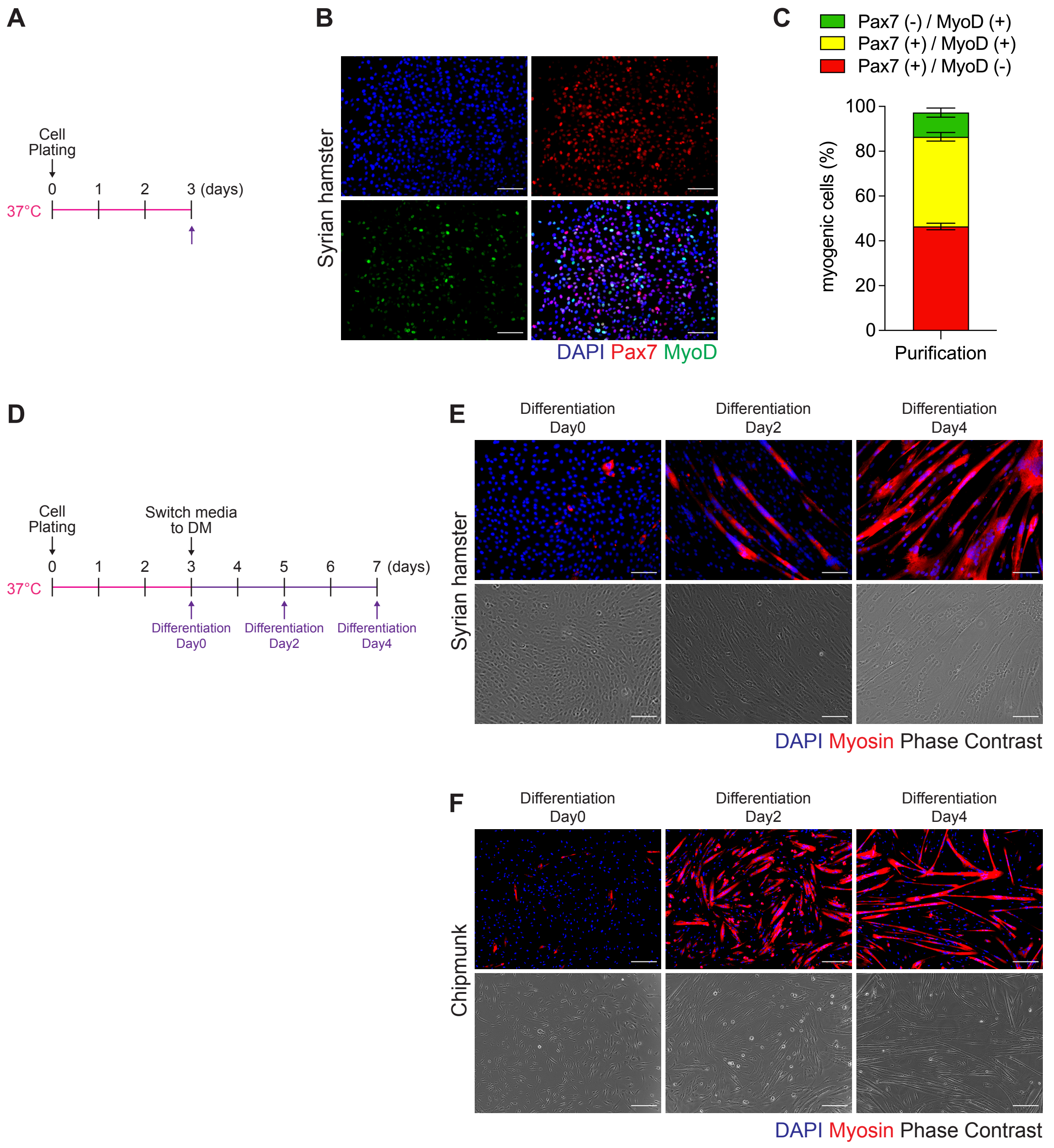


**Supplemental Figure 1 Characterization of satellite cells isolated from hibernating animals**

(A) Schematic overview of the experimental design for satellite cell (SC) isolation and characterization. SCs were plated immediately after isolation and maintained for 3 days at 37°C, with the growth medium replaced every other day. (B) Representative immunofluorescence images showing DAPI (blue), Pax7 (red), and MyoD (green) staining of Syrian hamster SCs. (C) Quantification of Pax7$^{+}$/MyoD$^{-}$, Pax7$^{+}$/MyoD$^{+}$, and Pax7$^{-}$/MyoD$^{+}$ cell populations. Each value represents the proportion relative to total cell number. Pax7- and/or MyoD-positive cells accounted for over 95% of the total, indicating high purity of the isolated SC population (n = 6). (D) Schematic overview of the experimental design for differentiation of SCs. Growth medium was switched to differentiation medium (DM) at day 3 post-plating, and DM was replaced every other day. (E) Representative images of Syrian hamster SCs induced to differentiate, showing DAPI (blue), myosin heavy chain (red), and phase contrast. (F) Representative images of chipmunk SCs after differentiation under the same conditions, demonstrating that their myogenic potential was preserved. Scale bars: 200 µm (B, E, F).
