## Supplemental_Figure_2 for "Cold-induced suppression of myogenesis in skeletal muscle stem cells contributes to delayed muscle regeneration during hibernation"

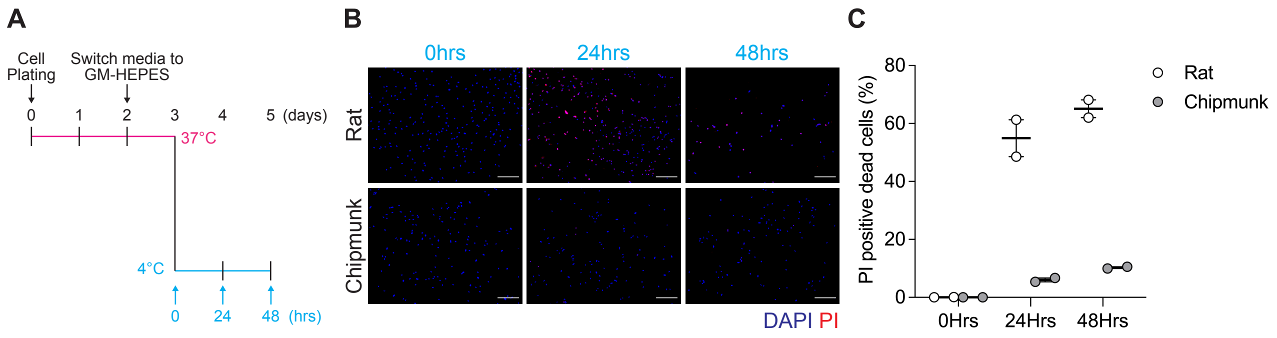


**Supplemental Figure 2 Cold-induced cell death in satellite cells from additional hibernating and nonhibernating species**

(A) Schematic overview of the experimental design to assess cold-induced cell death (CICD) in skeletal muscle satellite cells (SCs) from rats (nonhibernators) and chipmunks (hibernators). SCs were cultured at 37°C for 48 hours after plating, followed by a medium change to GM-HEPES and an additional 24-hour incubation at 37°C. Cells were then subjected to cold exposure at 4°C for 24 or 48 hours without changing the medium. (B) Representative fluorescence images showing DAPI (blue) and propidium iodide (PI; red) staining at each time point. (C) Quantification of PI-positive cells across time points. Rat SCs exhibited a time-dependent increase in PI-positive nuclei, whereas chipmunk SCs showed minimal changes. Data are presented as mean ± SD from two biological replicates per species. No statistical analyses were performed. Scale bars: 200 µm (B).
