## Supplemental_Figure_3 for "Cold-induced suppression of myogenesis in skeletal muscle stem cells contributes to delayed muscle regeneration during hibernation"

### Original western blotting images & Detailed experimental conditions
### The rectangle area with white dotted line was cropped and used for the figure.

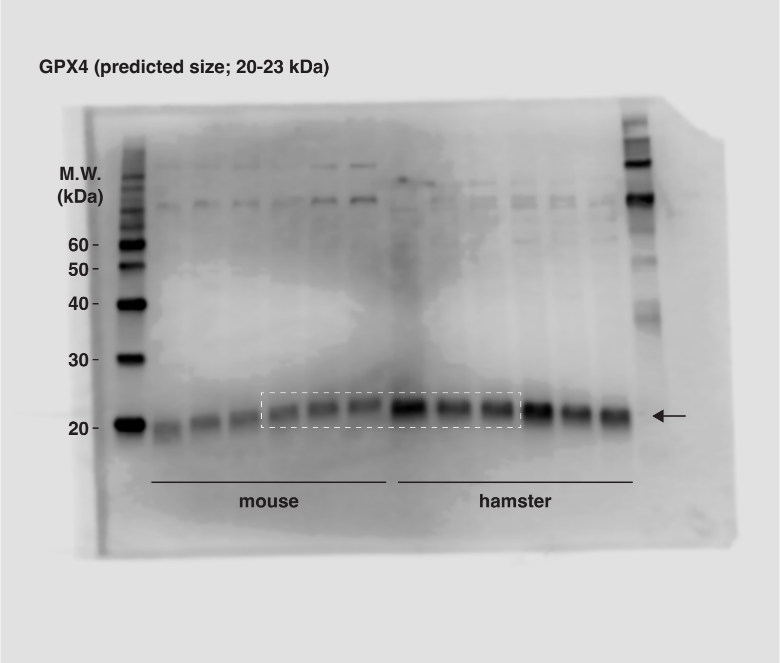


Protein: 300 $\mu$l of 1X sample buffer was added to a 35 mm dish and directly lysed
10 $\mu$l of sample was used for each lane

Gel system: TGX Any kD Precast Protein Gels, #4569036; BIO-RAD

Electrophoresis: 200V constant, 30-40 min

Transfer: Transblot Turbo; BIO-RAD, 2.5A contant, 3 min

Membrane: Trans-Blot TurboTM RTA Transfer PVDF Kit

Blocking: Bullet Blocking One, 5 min at r.t.

Primary Antibody: GPX4 Monoclonal antibody, rabbit
1:1000 dilution, 4°C, Over Night

2nd Antibody: Peroxidase Goat Anti-Rabbit IgG (H+L)
1:15000 dilution, 60 min at r.t.

Detection: ImmunoStar LD

Scan & Quantification: C-DiGit Blot Scanner

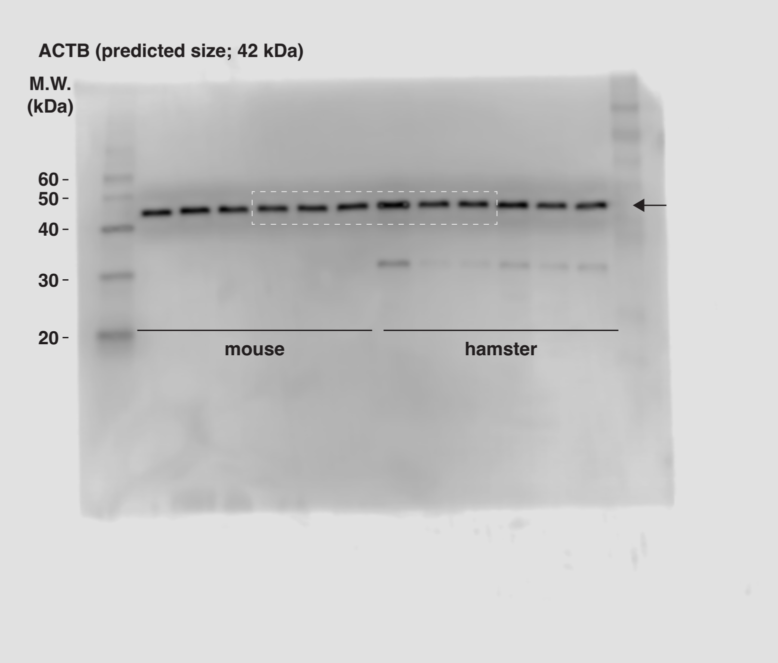


Protein: 300 $\mu$l of 1X sample buffer was added to a 35 mm dish and directly lysed
10 $\mu$l of sample was used for each lane

Gel system: TGX Any kD Precast Protein Gels, #4569036; BIO-RAD

Electrophoresis: 200V constant, 30-40 min

Transfer: Transblot Turbo; BIO-RAD, 2.5A contant, 3 min

Membrane: Trans-Blot TurboTM RTA Transfer PVDF Kit

Blocking: Bullet Blocking One, 5 min at r.t.

Primary Antibody: Beta Actin Monoclonal antibody, mouse
1:20000 dilution, 4°C, Over Night

2nd Antibody: Peroxidase Goat Anti-Mouse IgG (H+L)
1:15000 dilution, 60 min at r.t.

Detection: ImmunoStar Zeta

Scan & Quantification: C-DiGit Blot Scanner

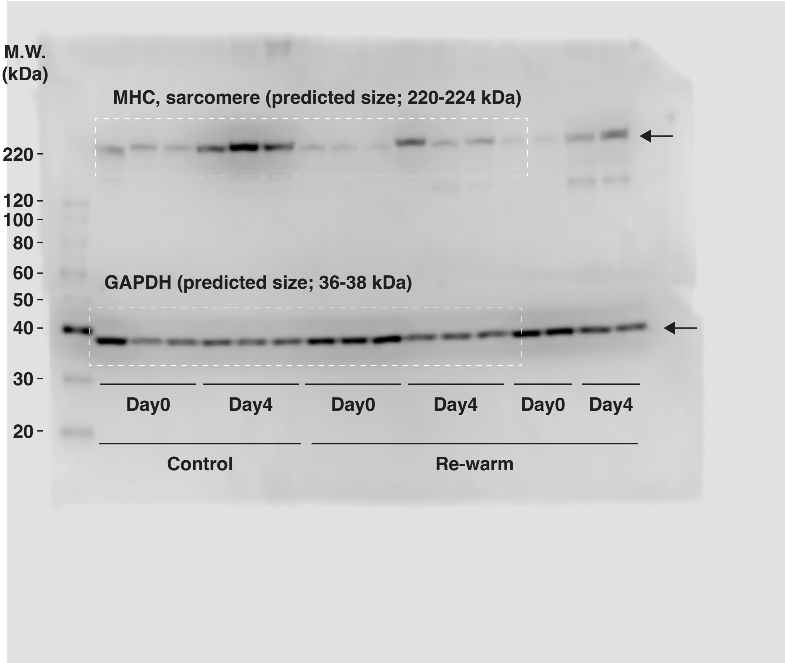


Protein: 300 μl of 1X sample buffer was added to a 35 mm dish and directly lysed
10 μl of sample was used for each lane

Gel system: E-R520L e-PAGEL (5-20% Gradient Gel)

Electrophoresis: 20mA constant, 75 min

Transfer: Criterion Blotter; BIO-RAD, 300mA contant, 90 min

Membrane: ClearTrans SP PVDF Membrane, 0.2μm

Blocking: Blocking One, 30 min at r.t.

Primary Antibody: MHC, mouse
0.2 ug/ml., 4°C, Over Night

GAPDH, mouse
1:5000 dilution, 4°C, Over Night

2nd Antibody: Peroxidase Goat Anti-Mouse IgG (H+L)
1:15000 dilution, 60 min at r.t.

Detection: ImmunoStar Zeta

Scan & Quantification: C-DiGit Blot Scanner

Protein samples were separated on the same gel and transferred onto a single PVDF membrane. After transfer, the membrane was cut horizontally between the 50 kDa and 60 kDa molecular weight markers to allow independent antibody probing for high- and low-molecular-weight proteins. The upper portion of the membrane was used for MHC detection, and the lower portion for GAPDH detection. Gel electrophoresis, membrane transfer, and blocking procedures were performed under identical conditions prior to membrane separation.

**Supplemental Figure 3 Original western blot images and detailed experimental conditions**

Original western blot images for glutathione peroxidase, β-actin, myosin heavy chain, and glyceraldehyde-3-phosphate dehydrogenase, along with detailed experimental conditions. Cropped areas indicated by white dotted lines in the original images were used in the corresponding main figures.
