## Supplemental_Table_1 for "Cold-induced suppression of myogenesis in skeletal muscle stem cells contributes to delayed muscle regeneration during hibernation"

| Category | Product/Kit Name | Manufacturer | Catalog No. | Application |
|---|---|---|---|---|
| Animal Care | Isoflurane Inhalation Solution | VIATRIS | 871119 | Anesthesia |
| Animal Care | Isoflurane | FUJIFILM Wako Pure Chemical Corporatio | 099-06571 | Anesthesia |
| Animal Care | Pentobarbital Sodium Salt | NACALAI TESQUE | 26427-72 | Anesthesia |
| Animal Care | Medetomidine Hydrochloride (Dorbene vet) | Kyoritsu Seiyaku Corp | QG03AA01 | Anesthesia |
| Animal Care | Midazolam (Midazolam Injection 10mg[SANDOZ]) | Sandoz K.K. | N05CD08 | Anesthesia |
| Animal Care | Butorphanol tartrate (Vetorphale) | Meiji Animal Health | N02AF01 | Anesthesia |
| Animal Care | Rodents Diets | Oriental Yeast | MF | Diets |
| Animal Care | Nosan Corporation Laboratory Animal Feed | Nihon Nosan | Labo MR Standa | Diets |
| Animal Care | Cardiotoxin | LATOXAN | L8102 | Muscle regeneration |
| Animal Care | iButton | Analog Devices | DS1922L-F5# | Temperature logger |
| Antibodies | Anti-Pax7 | Santa Cruz Biotechnology | sc-81648 | Immunofluorescence |
| Antibodies | Anti-MyoD | Santa Cruz Biotechnology | sc-760 | Immunofluorescence |
| Antibodies | Myog Antibody (F5D) | Developmental Studies Hybridoma Bank | F5D | Immunofluorescence |
| Antibodies | MYH1E Antibody (MF 20) | Developmental Studies Hybridoma Bank | MF 20 | Immunofluorescence |
| Antibodies | MYH3 Antibody (F1.652) | Developmental Studies Hybridoma Bank | F1.652 | Immunofluorescence |
| Antibodies | CoraLite Plus 488-conjugated CD68 Polyclonal antibody | Proteintech | CL488-28058 | Immunohistochemistry |
| Antibodies | CD206 Polyclonal antibody | Proteintech | 18704-1-AP | Immunohistochemistry |
| Antibodies | Anti-iNOS | FUJIFILM Wako Pure Chemical Corporatio | 015-28371 | Immunohistochemistry |
| Antibodies | Goat anti-Mouse IgG (H+L) Cross-Adsorbed Secondary Antibody, Alexa Fluor 594 | Thermo Fisher Scientific | A-11005 | Secondary antibody |
| Antibodies | Goat anti-Rabbit IgG (H+L) Cross-Adsorbed Secondary Antibody, Alexa Fluor 488 | Thermo Fisher Scientific | A-11008 | Secondary antibody |
| Antibodies | Goat anti-Mouse IgG1 Cross-Adsorbed Secondary Antibody, Alexa Fluor 488 | Thermo Fisher Scientific | A-21121 | Secondary antibody |
| Antibodies | Goat anti-Mouse IgG2b Cross-Adsorbed Secondary Antibody, Alexa Fluor 594 | Thermo Fisher Scientific | A-21145 | Secondary antibody |
| Antibodies | CoraLite594 - conjugated Goat Anti-Rabbit IgG(H+L) | Proteintech | SA00013-4 | Secondary antibody |
| Antibodies | Anti-Glutathione Peroxidase 4 antibody [EPNCIR144] | abcam | ab125066 | Western blot |
| Antibodies | Beta Actin Monoclonal antibody | Proteintech | 66009-1-Ig | Western blot |
| Antibodies | GAPDH (6C5) | Santa Cruz Biotechnology | sc-32233 | Western blot |
| Antibodies | Peroxidase AffiniPure Goat Anti-Rabbit IgG (H+L) | Jackson ImmunoResearch | 111-035-003 | Western blot |
| Antibodies | Peroxidase AffiniPure Goat Anti-Mouse IgG (H+L) | Jackson ImmunoResearch | 115-035-003 | Western blot |
| Instruments | Multiskan GO Microplate Reader | Thermo Fisher Scientific | 1510 | Absorbance measurement |
| Instruments | BZ-X800 Microscope System | Keyence | BZ-X800 | Imaging |
| Instruments | 5300 Fragment Analyzer System | Agilent Technologies | 5300 | RNA quality check |
| Instruments | DNBSEQ-G400 Sequencer | MGITech | DNBSEQ-G400 | RNA-seq |
| Instruments | Cryostat | Thermo Fisher Scientific | NX50 | Tissue sectioning |
| Instruments | Cryostat | Thermo Fisher Scientific | HM525NX | Tissue sectioning |
| Instruments | C-DiGit Blot Scanner | LI-COR Biosciences | 3600-00 | Western blot imaging |
| Reagents/Kits | FlexAble CoraLite Plus 488 Antibody Labeling Kit for Rabbit IgG | Proteintech | KFA001 | Antibody labeling |
| Reagents/Kits | FlexAble 2.0 CoraLite Plus 555 Antibody Labeling Kit for Rabbit IgG | Proteintech | KFA502 | Antibody labeling |
| Reagents/Kits | Z-VAD-FMK | Selleck | S7023 | Apoptosis inhibitor |
| Reagents/Kits | Penicillin-Streptomycin Solution (×100) | FUJIFILM Wako Pure Chemical Corporatio | 168-23191 | Cell culture |
| Reagents/Kits | D-MEM(High Glucose) with L-Glutamine, Phenol Red and Sodium Pyruvate | FUJIFILM Wako Pure Chemical Corporatio | 043-30085 | Cell culture |
| Reagents/Kits | -Cellstain- PI solution | Dojindo | P378 | Cell death assay |
| Reagents/Kits | -Cellstain- DAPI solution | Dojindo | D523 | Cell death assay |
| Reagents/Kits | 4%-Paraformaldehyde Phosphate Buffer Solution | NACALAI TESQUE | 09154-56 | Cell death assay |
| Reagents/Kits | Cytotoxicity LDH Assay Kit-WST | Dojindo | 343-91753 | Cytotoxicity assay |
| Reagents/Kits | FerroOrange | Dojindo | 342-09533 | Ferroptosis detection |
| Reagents/Kits | Ferrostatin-1 | Cayman Chemical | 17729 | Ferroptosis inhibitor |
| Reagents/Kits | Chick Embryo Extract, Ultrafiltrate, Liquid (CEE) | USBiological | C3999 | Growth medium |
| Reagents/Kits | Recombinant Murine FGF-basic | PeproTech | 450-33 | Growth medium |
| Reagents/Kits | Isopentane | NACALAI TESQUE | 26404-75 | Histology |
| Reagents/Kits | Mayer's hematoxylin solution | MUTO PURE CHEMICALS | 88591 | Histology |
| Reagents/Kits | 1% Eosin Y Solution | MUTO PURE CHEMICALS | 32002 | Histology |
| Reagents/Kits | Triton X-100 | NACALAI TESQUE | 35501-15 | Immunofluorescence |
| Reagents/Kits | DAPI-Fluoromount-G | SouthernBiotech | 0100-20 | Immunohistochemistry |
| Reagents/Kits | Necrostatin-1 | Cayman Chemical | 11658 | Necroptosis inhibitor |
| Reagents/Kits | Click-iT EdU Cell Proliferation Kit for Imaging, Alexa Fluor 488 dye | Thermo Fisher Scientific | C10337 | Proliferation assay |
| Reagents/Kits | ISOGEN II | NIPPON GENE | 311-07361 | RNA extraction |
| Reagents/Kits | RNeasy Mini Kit | Qiagen | 74104 | RNA extraction |
| Reagents/Kits | ROS Assay Kit -Highly Sensitive DCFH-DA | Dojindo | 340-09811 | ROS detection |
| Reagents/Kits | Collagenase Type II | Worthington Biochemical Corporation | CLS2 | SC isolation |
| Reagents/Kits | Matrigel | Corning | 354230 | SC isolation |
| Reagents/Kits | Injection Needle Pink 18G | Terumo | NN-1838R | SC isolation |
| Reagents/Kits | Cell Strainer | AS ONE | VCS-40 | SC isolation |
| Reagents/Kits | Collagen TypeI coated Dish 60mm | AGC TECHNO GLASS | 4010-010 | SC isolation |
| Reagents/Kits | Dimethyl Sulfoxide | NACALAI TESQUE | 13408-64 | Solvent |
| Reagents/Kits | MyJector Needle Implantable Type Insulin Syringe 29G | Terumo | SS-10M2913 | Western blot |
| Reagents/Kits | Sample Buffer Solution with 3-Mercapto-1,2-propanediol (×4) | FUJIFILM Wako Pure Chemical Corporatio | 196-16142 | Western blot |
| Reagents/Kits | Any kD Mini-PROTEAN TGX Precast Protein Gels | Bio-Rad Laboratories | 4569036 | Western blot |
| Reagents/Kits | Trans-Blot Turbo RTA Mini 0.2 µm PVDF Transfer Kit | Bio-Rad Laboratories | 1704272 | Western blot |
| Reagents/Kits | E-R520L e-PAGEL | ATTO Corporation | 2331730 | Western blot |
| Reagents/Kits | ClearTrans SP PVDF Membrane, Hydrophobic, 0.2 μm | FUJIFILM Wako Pure Chemical Corporatio | 033-22453 | Western blot |
| Reagents/Kits | Bullet Blocking One for Western Blotting | NACALAI TESQUE | 13779-01 | Western blot |
| Reagents/Kits | Blocking One | NACALAI TESQUE | 03953-95 | Western blot |
| Reagents/Kits | ImmunoStar LD | FUJIFILM Wako Pure Chemical Corporatio | 290-69904 | Western blot |
| Reagents/Kits | ImmunoStar Zeta | FUJIFILM Wako Pure Chemical Corporatio | 295-72404 | Western blot |

**Supplemental Table 1. List of materials, reagents, and equipment**

Detailed list of animal care materials, cell culture reagents, chemicals, staining reagents, antibodies, molecular biology kits, and other laboratory equipment. The table includes product names, manufacturers, catalog numbers, and specific applications as referenced in the Materials and Methods section.
