## Supplemental_Table_2 for "Cold-induced suppression of myogenesis in skeletal muscle stem cells contributes to delayed muscle regeneration during hibernation"

| Sample | Antigen | Primary Antibody | Secondary Antibody | Antibody Labeling Kit | Application |
|---|---|---|---|---|---|
| Hamster satellite cells | Pax7 | anti-Pax-7, mouse monoclonal, IgG1 kappa light chain | Goat anti-Mouse IgG (H+L), Alexa Fluor 594 | – | Immunofluorescence |
| Hamster satellite cells | MyoD | anti-MyoD, rabbit polyclonal, IgG | Goat anti-Rabbit IgG (H+L), Alexa Fluor 488 | – | Immunofluorescence |
| Hamster satellite cells | Myogenin | anti-Myogenin, mouse monoclonal, IgG1 kappa light chain | Goat anti-Mouse IgG (H+L), Alexa Fluor 594 | – | Immunofluorescence |
| Hamster satellite cells | Myosin heavy chain, sarcomere (MHC) | anti-Myosin heavy chain, mouse monoclonal, IgG2b, kappa light chain | Goat anti-Mouse IgG2b, Alexa Fluor 594 | – | Immunofluorescence |
| Hamster skeletal muscle tissue | Myosin heavy chain (embryonic) | anti-embryonicMHC, mouse monoclonal, IgG1 | Goat anti-Mouse IgG1, Alexa Fluor 488 | – | Immunohistochemistry |
| Chipmunk satellite cells | Myosin heavy chain, sarcomere (MHC) | anti-Myosin heavy chain, mouse monoclonal, IgG2b, kappa light chain | Goat anti-Mouse IgG2b, Alexa Fluor 594 | – | Immunofluorescence |
| Hamster skeletal muscle tissue | Pax7 | anti-Pax-7, mouse monoclonal, IgG1 kappa light chain | Goat anti-Mouse IgG1, Alexa Fluor 488 | – | Immunohistochemistry |
| Hamster skeletal muscle tissue | MyoD | anti-MyoD, rabbit polyclonal, IgG | Goat anti-Rabbit IgG (H+L), CoraLite594 | – | Immunohistochemistry |
| Hamster skeletal muscle tissue | CD68 | CoraLite Plus 488-conjugated CD68 Polyclonal antibody | – | – | Immunohistochemistry |
| Hamster skeletal muscle tissue | iNOS | anti-iNOS, rabbit polyclonal, IgG | – | FlexAble 2.0 CoraLite Plus 555 Antibody Labeling Kit for Rabbit IgG | Immunohistochemistry |
| Hamster skeletal muscle tissue | CD206 | anti-CD206, rabbit polyclonal, IgG | – | FlexAble 2.0 CoraLite Plus 555 Antibody Labeling Kit for Rabbit IgG | Immunohistochemistry |

**Supplemental Table 2. Antibodies and labeling kits used for immunofluorescence and immunohistochemistry**

List of primary and secondary antibodies used for immunofluorescence and immunohistochemistry, along with host species and corresponding antibody labeling kits. The table includes antigen targets, antibody specifications, and applications as referenced in the Materials and Methods section.
